# Multimodal Phasor Approach to study breast cancer cells invasion in 3D spheroid model

**DOI:** 10.1101/2024.06.10.598307

**Authors:** Giulia Tedeschi, Francesco Palomba, Lorenzo Scipioni, Michelle A. Digman

## Abstract

We implemented a multimodal set of functional imaging techniques optimized for deep-tissue imaging to investigate how cancer cells invade surrounding tissues and how their physiological properties change in the process. As a model for cancer invasion of the extracellular matrix, we created 3D spheroids from triple-negative breast cancer cells (MDA-MB-231) and non-tumorigenic breast epithelial cells (MCF-10A). We analyzed multiple hallmarks of cancer within the same spheroid by combining a number of imaging techniques, such as metabolic imaging of NADH by Fluorescence Lifetime Imaging Microscopy (NADH-FLIM), hyperspectral imaging of a solvatochromic lipophilic dye (Nile Red) and extracellular matrix imaging by Second Harmonic Generation (SHG). We included phasor-based bioimage analysis of spheroids at three different time points, tracking both morphological and biological properties, including cellular metabolism, fatty acids storage, and collagen organization. Employing this multimodal deep-imaging framework, we observed and quantified cancer cell plasticity in response to changes in the environment composition.

## Introduction

Crucial improvements have been made in understanding the development and progression of breast cancer, ranging from new methods for early diagnosis and to establishing new, more efficacious treatments ^1–3^. Despite this, breast cancer continues to be a leading cause of death, often associated with failure of critical organs due to metastasis. The ability of cancer cells to invade the surrounding tissue and the bloodstream is the first step towards the formation of metastatic tumors ^4^. The local invasion occurs through the basement membrane layer, composed mainly of collagen IV and laminins, and then through the dense extracellular matrix constituted in large part by fibrils of collagen I ^5,6^. In recent years, many studies have acknowledged the importance of a 3D in vitro cell model system to mimic in controlled conditions the complexities of an in vivo solid tumor and its microenvironment. 2D monolayer culture often fails to predict drug efficiencies as well as migratory and invading properties of breast cancer cells. Three-dimensional multicellular tumor spheroids, a controlled 3D culture of cancer cells, recapitulates several architectural and biological hallmarks and represent a valid bridge to cover the gap between the simplified 2D monolayer and the complexity of solid tumors in animal models ^7–10^. Once embedded in a substrate that mimics the extracellular matrix (ECM), this 3D model system can also provide valuable information about the cell-cell interaction inside the tumor as well as cell-extracellular matrix mechanical and biochemical interactions ^11^. For instance, spheroids created from the MDA-MB-231 triple negative breast cancer cell line showed an upregulation of EMT-associated proteins, different migratory and proliferation properties and increased resistance to antitumor compounds that couldn’t be addressed with 2D culture ^12^. Therefore, in vitro tumor spheroids represent a valuable platform not only to give insights into the processes that lead to the ECM invasion but also to evaluate the efficacy of drug treatments and to improve drug development studies ^13–15^.

Numerous studies have supported the concept that cancer development and progression are driven by the cross-talk between cancer cells and the surrounding microenvironment ^16,17^. A striking example of this strong connection is the link between poor patient survival and the increase in fiber density accompanied with formation of bundled aligned collagen surrounding the tumor ^18^. ECM provides structural and mechanical support to cells and contributes to the regulation of gene expression, cell cycle, apoptosis and migration ^19^. One of the earliest steps of breast cancer metastasis is cellular migration and invasion through the surrounding collagenous stromal extracellular matrix. A primary tumor subset of cells termed “leaders’’ start to degrade and remodel the surrounding matrix secreting proteolytic enzymes: this creates invasion paths (micro tracks) that “follower” cells can later use to migrate out the primary tumor, without need of proteolytic enzymes ^20^. Several ECM components, including fibrillar collagens, fibronectin, laminins and hyaluronic acid, as well as ECM remodeling enzymes are specifically induced and play a major functional role in breast cancer progression ^21–23^.

Cell motility and their interaction with the ECM has also been correlated with metabolic changes in several different systems ^24–27^. One of the hallmarks of cancer is the concept of a metabolic switch to glycolysis (Warburg effect ^28^), where other energy pathways (i.e. Krebs cycle and oxidative phosphorylation) are frequently shut down or reduced. However, recent studies have highlighted considerable metabolic heterogeneity of cancer cells inside solid tumors, their ability to switch and adapt their metabolism during tumor progression ^26,29–32^. For instance, in some types of cancer cells oxidative phosphorylation metabolism is required to migrate ^33^. Metabolic plasticity of breast cancer cells has also been associated with switches between active and quiescent states ^34^. Understanding the consequences of changes in metabolism to tumor progression can provide useful insight for cancer treatments.^35^

In terms of lipid metabolism, intracellular lipid droplets have been shown to play a fundamental role in the cross-talk between cancer cells and the tumor microenvironment ^36–39^. In particular, lipid droplets not only act as a reserve of energy but they also contribute to the cellular stress adaptive response ^40,41^, protecting the cells against hypoxia ^42,43^, lipotoxicity and oxidative stress. Lipid droplets can regulate cell metabolism, proliferation, migration and invasiveness ^44–46^ and, through their complex functions, they can contribute to tumor progression ^47^ and resistance to chemotherapeutic treatments ^48,49^.

These three cancer relevant hallmarks (metabolic switch, lipid metabolism and collagen remodeling) have never been observed simultaneously in invading tumor spheroids due to challenging multiplex feasibility and microscopy techniques required. Here, we present a framework for the acquisition, on the same spheroid, at single cell level, of metabolic state, lipid composition and collagen remodeling by combining FLIM-NADH imaging, lipid order/composition through spectral imaging of a solvatochromic dye and extracellular matrix morphology and structure at nanoscale using second harmonic generation (SHG) imaging.

Tumor spheroids are embedded in collagen to mimic the interaction with the extracellular matrix (ECM). This configuration poses challenges as the optical penetration in the sample is limited, requiring deep-tissue imaging approaches to ensure quantification of the spheroid in its entirety. To do so, we turned to 2-photon microscopy for metabolic imaging and second harmonic generation all the while using a lipophilic dye, Nile Red, to ensure labeling of the deeper layers of the spheroid while reporting on lipid composition. In order to monitor the ECM modifications carried out by invasive cells, we use SHG imaging, collecting the signal produced by the light scattered from the highly ordered fibers of the biopolymers in the ECM, combined with PLICS (Phasor Local Image Correlation Spectroscopy) ^50^, an image-based correlation analysis that allows to map pixel-wise the statistical distribution of the fiber size and density. Fluorescence Lifetime Imaging Microscopy ^51^ (FLIM), combined with the phasor approach ^52,53^ utilize the free-to-bound ratio of the coenzyme nicotinamide adenine dinucleotide (NADH) to quantify the balance between glycolysis (GLY) and oxidative phosphorylation (OXPHOS). When NADH is bound to enzymes, it has a longer lifetime (τ∼3.2 ns), while when it is free it has a shorter lifetime (τ∼0.4 ns). NADH-FLIM signature has been correlated in many previous studies to cellular metabolism ^54–57^. Finally, we made use of Nile Red, a solvatochromic dyes that exhibits a red-shift of the spectral emission that depends on the polarity of the solvent ^58^. In cells, Nile Red preferentially accumulates in lipid droplets, as they are highly hydrophobic/apolar, and its spectral emission can be analyzed with spectral phasors to obtain a parameter that correlates with lipid storage ^59^. Furthermore, the hydrophobicity of Nile Red is advantageous as it allows the uniform labeling of the spheroids, including their interior. In this work, we use this framework to investigate how these pathways of interest change between aggressive breast cancer cells (MDA-MB-231) and normal breast epithelial cells (MCF10a) in culturing conditions and in response to decreased collagen stiffness, glucose deprivation and oleic acid treatment.

## Results

### Multimodal approach enables the capture of metabolism, lipid profile and collagen structures in the same spheroid at different time points

To model the first phases of cancer cells leaving the main tumor mass to invade the surrounding matrix, we created a 3D spheroid grown from a human triple negative breast cancer immortalized cell line, MDA-MB-231, and a non-tumorigenic breast epithelial cell line, MCF-10A. After creating a number of spheroids in a 96-well plate (see material and methods), we embedded each spheroid in a 3D collagen Type I matrix and followed the spheroid evolving over 24 hours (**Figure 1A**). During this time, in triple negative breast cancer spheroids, some cells leave the main body and invade the matrix, we will refer to this subpopulation as “invasions”. To capture as many cancer hallmarks as possible, we adopted a multimodal approach capable of monitoring, in parallel and in the same spheroid, morphological properties (e.g., distance of the perimeter of the spheroids from the center) as well as properties related to cellular metabolism, lipid profile and collagen organization. At each time point considered (2 hours, 12 hours and 24 hours after collagen embedding), we carried out sequentially three different types of measures, as shown in **Figure 1**. We measured the lifetime of NADH autofluorescence (FLIM-NADH, **Figure 1B, left**) to determine the ratio between free NADH and enzyme-bound NADH. This ratio relates to whether cellular metabolism is relying more on glycolysis (low ratio) or oxidative phosphorylation (high ratio). To evaluate lipid profile, we measured the spectral emission of Nile Red (**Figure 1B, center**), an environmental sensitive dye that features a red-shift in its emission spectrum when interacting with highly polar lipids (such as phospholipids in the plasma membrane and internal membranes) and a blue-shift when interacting with environments with a lower polarity (such as lipid droplets). Finally, we used Second Harmonic Generation (SHG) to visualize the collagen matrix fibers (**Figure 1B, right**). For each technique applied we developed a phasor-based analysis pipeline. In particular we used lifetime phasors to evaluate the metabolic shift based on NADH autofluorescence (**Figure 1C, left**), spectral phasors to measure differences in emission of Nile Red (**Figure 1C, center**) and PLICS phasors to measure properties of the collagen fibers (**Figure 1C, right**).

**Figure 1.**
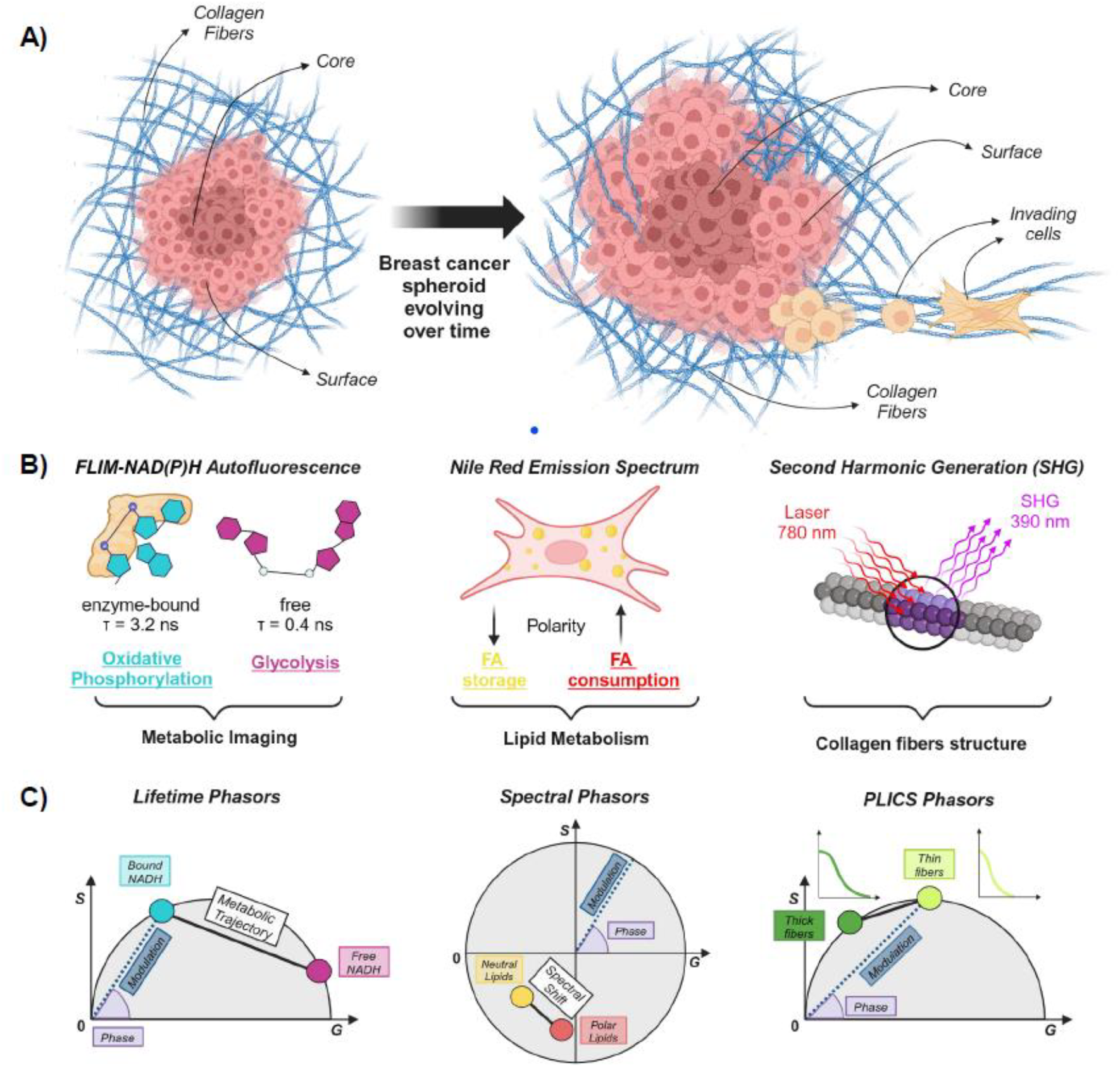
Schematic representation of the multimodal image acquisition, data analysis and spheroids evolution over time. **(A)** Left, a breast cancer spheroid embedded in a 3D collagen matrix at the first time point considered and how it evolves over 24 hours time (right), when some cells leave the main body to invade the surrounding collagen matrix. **(B)** Left to right, the measurement of lifetime of NADH autofluorescence for metabolic imaging, the emission spectral shift of the environmental sensitive dye Nile Red to measure lipid storage, the use of Second Harmonic Generation to visualize collagen fibers. **(C)** Left to right, Lifetime Phasors to measure the metabolic trajectory and the fraction of bound NADH, Spectral Phasors to measure the shift in the emission spectrum of Nile red depending of the lipid environment conditions, PLICS Phasors to quantify the thickness of collagen fibers measured with the SHG.

Our first aim in order to achieve an optimized multimodal system to study the evolution of 3D cancer spheroids over time was to make sure that the presence of Nile Red as a lipid metabolism probes did not interfere with measurements of NADH autofluorescence.

As shown in **Figure S1**, we performed FLIM-NADH measurements using a 2-Photon laser excitation of 720nm, optimizing the NADH excitation while being well separated from the excitation spectra of Nile Red. Secondly, we introduced a 419/46 emission filter for collecting photons in a spectral window that rejects potential bleed through from Nile Red fluorescence. For collection of the Nile Red spectral signal, we tuned the 2-Photon excitation laser at 800 nm, minimizing NADH excitation. Using these parameters, we proved that Nile Red presence does not affect the FLIM NADH readout by comparing unlabeled controls to Nile Red treated samples, both in single cells embedded in collagen and in 3D spheroids, as shown in **Figure S2**. Next, we tested different focal planes in our 3D spheroid model, using as a reference the bottom imaging plane of each spheroid, ensuring the acquisition of the representative section with optimal signal in FLIM-NADH, Nile Red spectral emission and SHG, as shown in **Figure S3**. As a result, to ensure consistent reproducible imaging, we chose as the optimal Z-plane the one located 100 μm from the bottom of the spheroid. After optimizing the imaging parameters, we moved to imaging live spheroids generated from MCF-10a and MDA-MB-231 cell lines.

### MCF-10a and MDA-MB-231 spheroids show unique hallmarks

Previous studies conducted on single cells in a 2D environment showed a metabolic difference measured with FLIM-NADH between MDA-MB-231 (human triple negative breast cancer cells) and MCF10a (non-tumorigenic human breast epithelial cells) ^5,6^. As shown in **Figure S4**, we observed a different response in terms of Nile Red spectral emission in 2D-cultured MDA-MB-231 cells and MCF10a cells. MDA-MB-231 cells show a shift of the Nile Red spectral emission towards shorter wavelength, denoting an enrichment for MDA-MB-231 cells in neutral lipids accumulated in lipid droplets compared to the contribution of the polar membrane phospholipids.

After assessing that metabolic and lipid profile differences between MDA-MB-231 and MCF10a can be detected in single cells in 2D cultures with both FLIM-NADH and Nile Red spectral emission, we proceeded to use these tools in a biologically more significant 3D spheroid model.

Spheroids grown from non-tumorigenic MCF10a cells and triple negative breast cancer cells MDA-MB231 were embedded in a 3D collagen I matrix, as described in the Methods section, and imaged after 2-, 12- and 24-hours post-embedding. Spheroids were segmented in two macro-regions, the core (at the interior of the spheroid) and the surface (in contact with the culturing medium), plus an additional category for cells invading the collagen matrix, which we found unique to MDA-MB-231 spheroids and detectable only at the 12- and 24-hour timepoints. In **Figure 2**, we report the multimodal profiling of MCF10A and MDA-MB-231 spheroids from the morphological (**Figure 2A**), NADH (**Figure 2B**) or lipid (**Figure 2C**) metabolism and collagen remodeling (**Figure 2D**) points of view. Comparing the spheroids from the two different cell lines we could observe that overall MDA-MB-231 spheroids show unique hallmarks compared to MCF10a spheroids, as we can appreciate from the representative images in **Figure 2E**. In particular, MDA-MB-231 spheroids show an increase in the distance between the perimeter and the center of the spheroid (**Figure 2A**), due to cells leaving the main tumor mass to invade the surrounding collagen. Moreover, MDA-MB-231 spheroids have a lower fraction of bound NADH compared to MCF10a spheroids, therefore relying on a more glycolytic metabolism compared to MCF10a (**Figure 2B**). Interestingly, we can observe a gradient of increased fraction bound as a function of time over the 24 hours as well as a function of the distance from the center of the spheroid. The cancer spheroids also show a shorter wavelength spectral emission of Nile Red, meaning an overall lower lipid polarity compared to the non-tumorigenic control (**Figure 2C**), indicating a higher storage of neutral lipids. We can also appreciate differences in the spatial organization of these properties, as MDA-MB-231 spheroids decrease their lipid polarity as a function of time and distance from the center at 2 hours, while at 24 hours the core is the compartment with lower polarity. From **Figure 2D** we can observe how MDA-MB231 spheroids are reshaping the collagen more than MCF10a spheroids, due to the cells migrating and invading the surrounding collagen (“inv”) and the contact between the cells at the interface between the main tumor mass and the collagen matrix (“surf”). We also observed a change in the collagen structure at 24 hours even in the regions far (“out”) from the MDA-MB231 spheroid body, likely due to the effect of some of the far-migrating cells. These results set a baseline for investigating lipid and NADH metabolism in tumor spheroids together with their collagen-reshaping capabilities. In the following, we will explore more in detail the effect that transient changes in environmental conditions have on triple-negative tumor spheroids.

**Figure 2.**
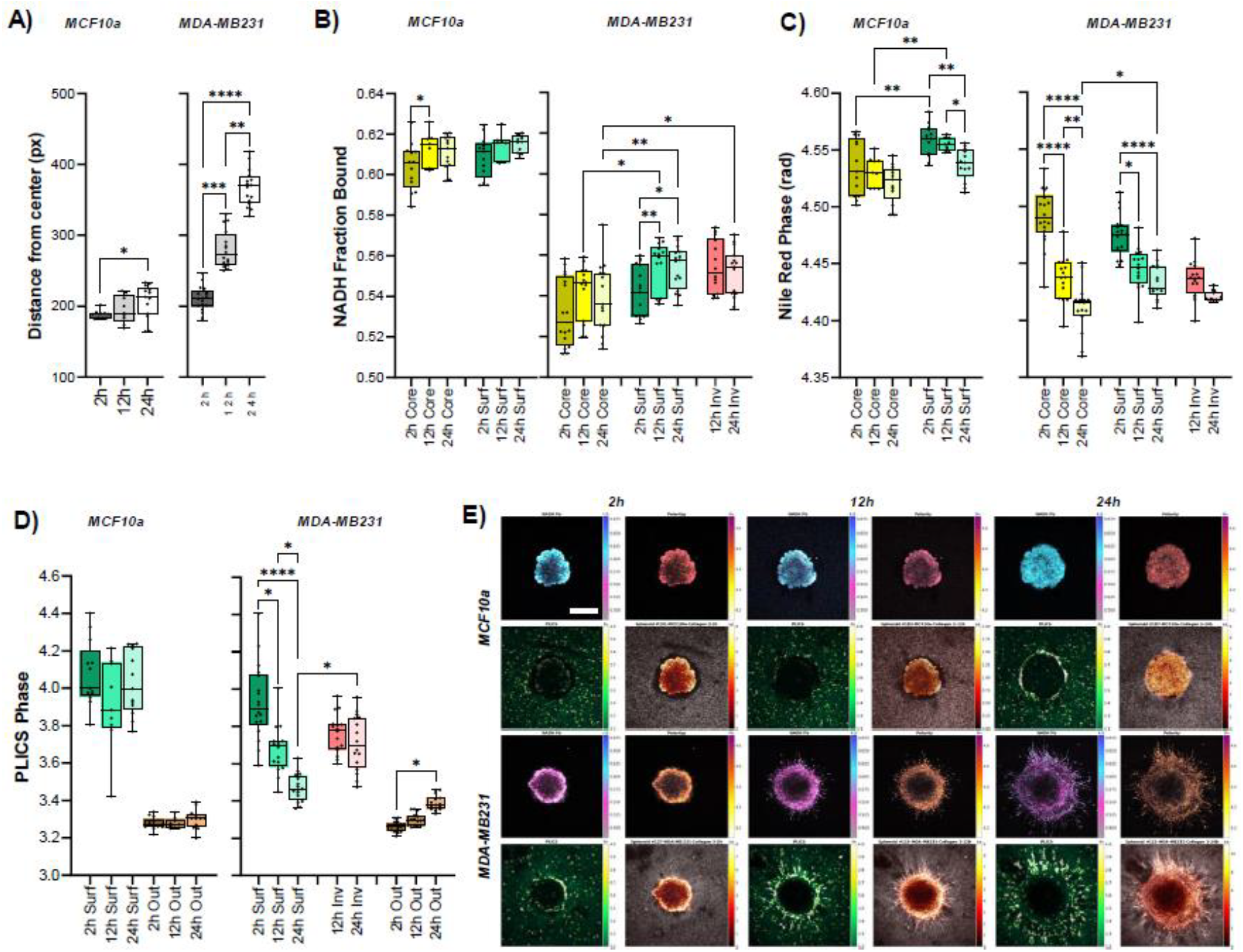
MCF-10a and MDA-MB-231 spheroids show unique hallmarks. **(A)** Distance of the perimeter of the spheroids from the center expressed in pixels as a function of time and cell line (MCF·Ioa, left; MDA-MB231, right). Metabolic shift measured as the fraction of Bound NADH **(B)** and Nile Red emission spectral shift **(C)** for different parts of the spheroids (Core, Surf = surface, Inv = cells invading the ECM) as a function of time and cell line (MCF10a, left; MDA-MB231, right). **(D)** PLICS Phase calculated for the collagen fibers in different regions (Surf = close to the surface of the spheroids, Inv = close to the invading cells, Out = far from the main body/invasions of the spheroids) as a function of time and cell line (MCF10a, left; MDA-MB231, right). Data are represented by box and whiskers plots. The box represent the median (solid line) and standard deviation. The whiskers go down to the smallest value and up to the largest All the points are shown. One-way ANOVA (K-W test), p>0 05 (ns), 0.05>p>0.01 (*), 0.01>p>0.001 (**). 0.001>p>0.0001 (***), p<0.0001 (****). **(E)** Representative images of Spheroids hallmarks (NADH Fraction Bound, Nile Red spectral emission, collagen PLICS, Nile Red and Collagen intensities) as a function of time and cell lines (MCF1 0a, top; MDA-MB231, bottom). Scale bar is 400μm.

### Effect of transient metabolic stress on MB231 spheroids

Next, we focused on MDA-MB-231 spheroids and, in particular, their recovery after transient stress. We exposed the spheroids to media without glucose (glucose starvation or GS) or supplemented exogenous fatty acids (oleic acid or OA) for 24 hours while the spheroids were growing in liquid culture. After the embedding in the collagen matrix, we supplemented the spheroids back with their normal growth medium to study the effect and the recovery after transient metabolic stress on collagen invasion and metabolism. The effect of the transient stress on metabolism, lipids, collagen remodeling and morphology is shown in **Figure 3**, where the data are represented as differences from the control (MDA-MB-231 spheroids in growth condition). Representative images of the effect of the transient stress on MDA-MB-231 spheroids over 24 hours after removing the stressing factor can be appreciated in **Figure 3A-B**.

**Figure 3.**
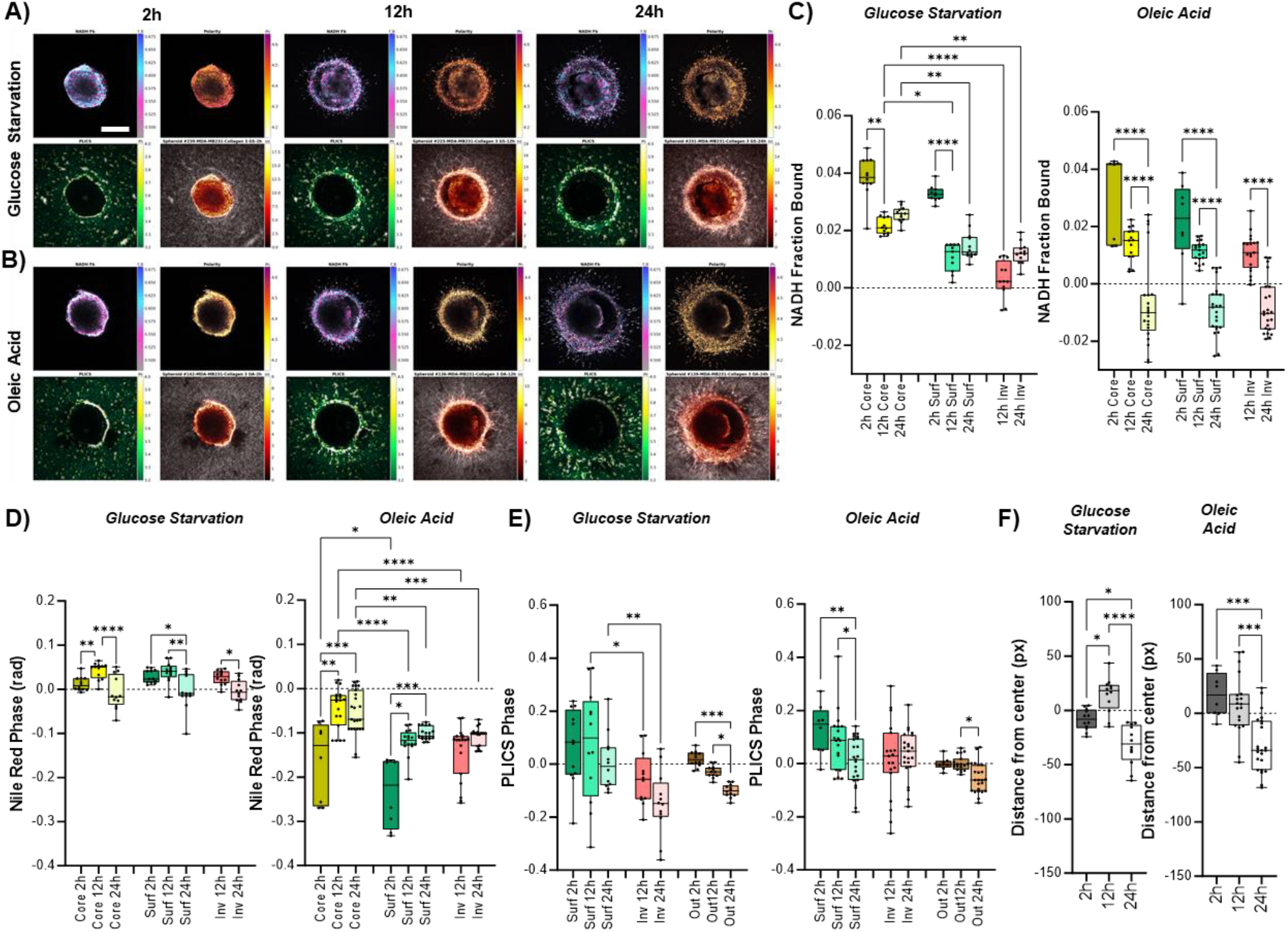
Effect of transient metabolic stress on MDA-MB231 spheroids. Representative images of MDA-MB231 spheroids hallmarks (NADH Fraction Bound, Nile Red spectral emission, collagen PLICS, Nile Red and Collagen intensities) as a function of time and treatments (Glucose Starvation, **(A)**; Oleic Acid, **(B)**). Scale bar is 400μm. Metabolic shift measured as the fraction of Bound NADH **(C)** and Nile Red emission spectral shift **(D)** for different parts of the spheroids (Core, Surf = surface, Inv = cells invading the ECM) as a function of time and treatment (Glucose Starvation, left; Oleic Acid, right). **(E)** PLICS Phase calculated for the collagen fibers in different regions (Surf= close to the surface of the spheroids, Inv= close to the invading cells, Out= far from the main body/invasions of the spheroids) as a function of time and treatments (Glucose Starvation, left; Oleic Acid, right). **(F)** Distance of the surface of the spheroids from the center expressed in pixels as a function of time and treatments (Glucose Starvation, left; Oleic Acid, right). All data in panels C-F are shown as difference from the mean value of the control, represented by the dashed black line. Data are represented by box and whiskers plots. The box represent the median (solid line) and standard deviation. The whiskers go down to the smallest value and up to the largest. All the points are shown. One-way ANOVA (K-W test), p>0.05 (ns), 0.05>p>0.01 (*), 0.01 >p>0.001 (**), 0.001>p>0.0001 (***), p<0.0001 (****).

In **Figure 3C** we can observe, during the first time point (2 hours), how both GS and OA treatments are shifting the balance between free and bound NADH towards the bound form, indicating a preference towards more OxPhos metabolism in the core and surface of the spheroids which persists at 24 hours, with cells at the surface and the cells invading the collagen shifting closer to the control while still maintaining an offset. Interestingly, with the OA treatment, core, surface and invasions follow the same metabolic trend, up until they all switch to a more glycolytic behavior at the 24-hour time point.

Regarding lipid storage, we can appreciate in **Figure 3D** how GS treatment induces a small shift in the Nile Red emission spectrum and for every macro-area of the spheroids, shift that is normalized back to values similar to the control in 24 hours. On the contrary, OA treatment at the first time point produces a huge shift towards lower lipid polarity and more neutral lipids content in all compartments. While this behavior is expected since OA is a precursor of triglycerides, we can appreciate a differential response for different macro-areas of the spheroid. Indeed, while the core reaches values closer to the control at 24 hours, surface and invasions partially reduce the initial shift but maintain a higher neutral lipid content compared to the control even at 24 hours. As this is not observed in MCF 10A spheroids (**Figure S5**) that completely revert back to control conditions, we are looking at a behavior specific to MDA-MB-231, that tends to maintain a larger storage of neutral lipids without consuming it. This observation, together with the sudden glycolytic switch at 24 hours, shows that even transient exposure to a lipid-rich environment can have long-term effects on metabolic properties. Furthermore, lipid droplets have been shown to be involved in mechanisms of chemoresistance and breast cancer stemness ^47^.

How do these metabolic changes translate in terms of collagen reshaping? As we can observe in **Figure 3E**, the GS treatment induces a higher collagen reshaping compared to the control, especially in the cells invading the matrix and in the collagen further from the main tumor body: in these regions we observe a decrease the thickness of the collagen fibers over 24 hours. Interestingly, OA treatment induces less collagen reshaping compared to GS and we can measure values similar to the control for the core at 24 hours, therefore the collagen-reshaping capability is not altered by the previously described metabolic changes, as can also be appreciated from the distance of the spheroid perimeter from the center, described in **Figure 3F**. Notably, with our multimodal approach we can appreciate the response not only of the surface of the spheroids and invading cells but also of the inner layers, usually less accessible for imaging.

### ECM density affects the metabolism and growth of MDA-MB231 spheroids

Finally, we investigate the effect of cells-ECM interactions by embedding the spheroids in a matrix with lower collagen density (1.5mg/mL compared to our control 3.0mg/mL). Representative images and relative properties of MDA-MB231 spheroids growing in less dense matrices at different time points are shown in **Figure 4A**. The first effect that we can appreciate in **Figure 4B** is a decrease of the average distance of the perimeter of the spheroids compared to the control, that become more pronounced at the 12- and 24-hour time points, indicating that the collagen-invading cells travel less far at lower collagen density. From the metabolic point of view, the analysis of FLIM-NADH measures in **Figure 4C** show an increase of the NADH fraction bound at 12 hours (indicating a shift towards ox-phos) followed by a decrease at 24 hours (indicating a shift towards glycolysis). We observe a similar trend in core, surface and invasions. The analysis of Nile Red emission spectrum shown in **Figure 4D** reveals a small change in the lipid profile, spatially confined at the cored, with no appreciable effect on the outer macro-areas of the spheroid. Interestingly, we can observe that MDA-MB-231 spheroids generate thicker collagen fibers at lower collagen density than they do in higher collagen density (**Figure 4E**), and that this effort does not alter the lipid storage of the spheroid, while it heavily relies on its NADH metabolic plasticity. Note that none of the changes observed for MDA-MB-231 spheroids applies to non-tumorigenic MCF10A spheroids that display no appreciable difference at any time point considered (**Figure S6**). Taken together, our multimodal analysis of cancer hallmarks can identify the metabolic profile of tumor spheroids, spatially and temporally resolved, and address their plasticity in diverse environmental conditions.

**Figure 4.**
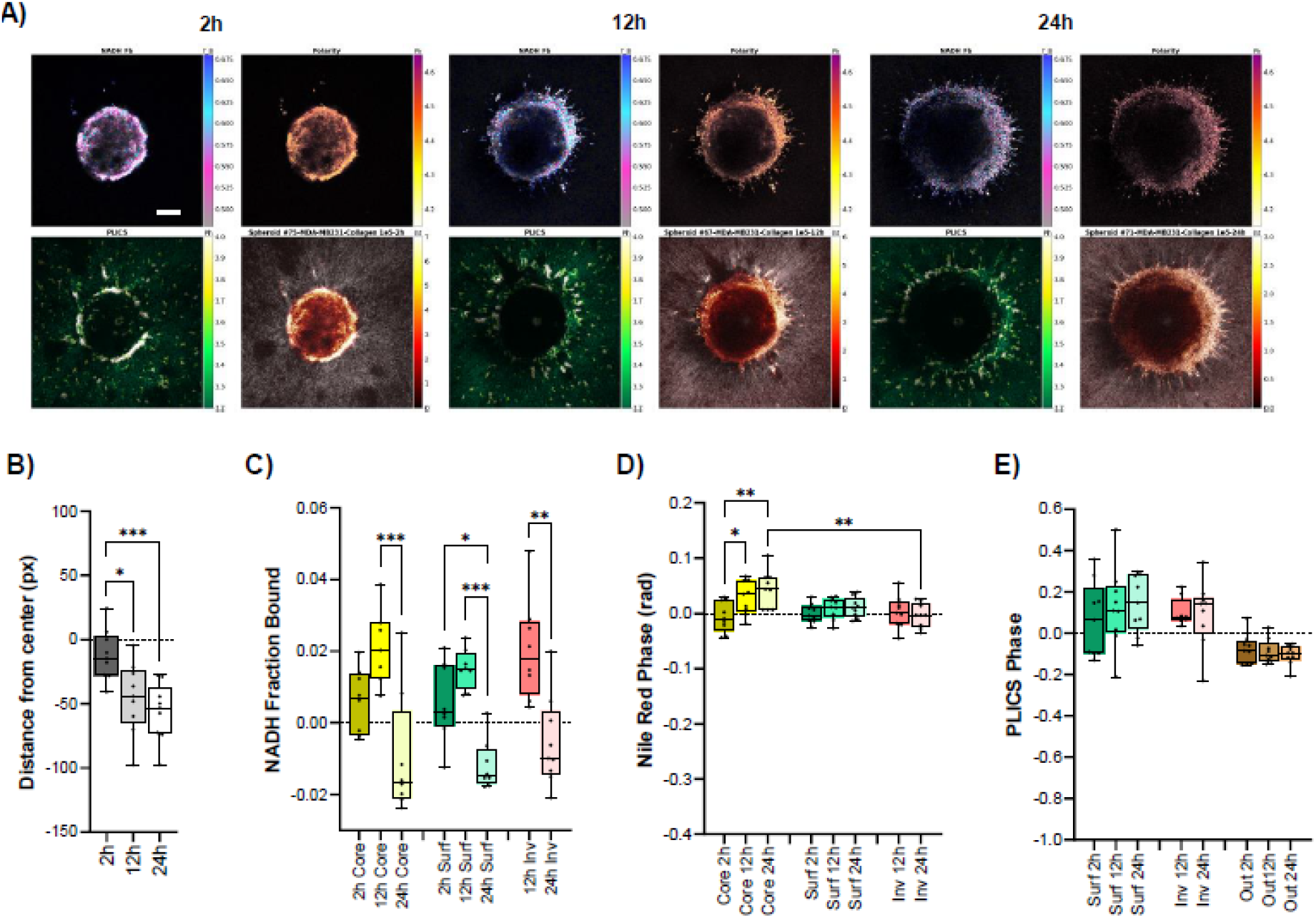
Effect of collagen density on MDA-MB231 spheroids. **(A)** Representative images of MDA-MB231 spheroids hallmarks (NADH Fraction Bound, Nile Red spectral emission, collagen PLICS, Nile Red and Collagen intensities) as a function of time when embedded in a collagen matrix with lower density Scale bar is 4l00μm **(B)** Distance of the surface of the spheroids from the center expressed in pixels as a function of time. Metabolic shift measured as the fraction of Bound NADH **(C)** and Nile Red emission spectral shift **(D)** for different parts of the spheroids (Core, Surf = surface, Inv = cells invading the ECM) as a function of time. **(E)** PLICS Phase calculated for the collagen fibers in different regions (Surf = close to the surface of the spheroids, Inv = close to the invading cells, Out = far from the main body/invasions of the spheroids) as a function of time. All data in panels **B-E** are shown as difference from the mean value of the control, represented by the dashed black line. Data are represented by box and whiskers plots The box represent the median (solid line) and standard deviation. The whiskers go down to the smallest value and up to the largest. All the points are shown. One-way ANOVA (K-W test), p>0.05 (ns), 0.05>p>0 01 (*), 0.01 >p>0.001 (**), 0 001 >p>0.0001 (***), p<0 0001 (****).

## Discussion

This study employed a deep-tissue multimodal imaging approach to investigate how breast cancer cells (MDA-MB-231) invade surrounding tissues and how their hallmarks change during this process. We utilized 3D spheroids embedded in a collagen matrix to mimic the tumor microenvironment and applied a cohort of techniques, including FLIM-NADH, hyperspectral imaging, and SHG to monitor metabolic state, lipid storage, and collagen remodeling, respectively. Our findings revealed distinct characteristics between cancer and non-tumorigenic (MCF-10A) cells and a distinct metabolic difference between the spheroid inner layers and invading cells, as observed in other studies ^30,59,60^. Furthermore, the cancer cells displayed remarkable metabolic plasticity, adapting their metabolic state and collagen remodeling activity in response to changes in the environment, such as glucose starvation, fatty acid exposure, and lower collagen density, maintaining their modified phenotype for extended periods of time. This adaptability highlights the complexity of cancer cell behavior and the importance of considering the 3D tumor microenvironment when developing therapeutic strategies, as the physiological properties of MDA-MB-231 cells are not only dependent on environmental conditions, but also to transient variation of its composition.

Future studies could explore the specific signaling pathways underlying the observed metabolic plasticity and collagen remodeling in cancer cells, highlighting the mechanisms that play a role in cancer plasticity depending on environmental cues. Additionally, investigating the effects of therapeutic agents targeting these pathways on cancer cell invasion in 3D models would be crucial for further development and translation into clinical/diagnostic applications. Furthermore, incorporating orthogonal techniques, such as gene or protein expression profiling, could provide even deeper insights into the complex interplay between metabolism, lipid storage, and the invasive behavior of cancer cells. Our study employed a single cancer cell line (MDA-MB-231) and a single non-tumorigenic cell line (MCF-10A). Investigating a broader range of cell lines with varying metastatic potentials could provide a more comprehensive understanding of the observed phenomena. Additionally, while our 3D spheroid model offers advantages over traditional 2D cultures, it still represents a simplified version of the complex tumor microenvironment. Utilizing more intricate models incorporating stromal cells and immune cells could provide a more realistic picture of cancer cell invasion in vivo. In conclusion, this study demonstrates the power of a multimodal imaging approach to investigate cancer cell invasion in a 3D microenvironment. The observed link between altered metabolism, increased neutral lipid storage, and enhanced collagen remodeling suggests a coordinated effort by cancer cells to invade surrounding tissues. Furthermore, the ability of these cells to switch metabolic states and remodel collagen in response to environmental changes highlights their remarkable plasticity. These findings not only provide new insights into cancer cell invasion but also offer potential targets for therapeutic intervention. Future studies could utilize our multimodal imaging platform to explore how targeting these metabolic pathways or the ability of cancer cells to remodel the collagen matrix might hinder their invasive behavior and metastatic spread.

## Material and Methods

### Cell Culture and 3D spheroid models

MDA-MB231 cells were cultured in DMEM medium (high glucose, with L-glutamine, GenClone), supplemented with 10% v/v Fetal Bovine Serum (heat inactivated, GenClone) and 1% v/v penicillin/streptomycin solution 100× (10,000 units of penicillin and 10 mg ml−1 streptomycin in 0.85% saline solution, GenClone), in a 37 °C and 5% CO2 incubator.

MCF10a cells were cultured in DMEM/F12 medium (Gibco), supplemented with 5% v/v Horse Serum (heat inactivated, Invitrogen), Epidermal Growth Factor (Peprotech), Hydrocortisone (Sigma), Cholera Toxin (Sigma), Insulin from bovine pancreas(Sigma), 1% v/v penicillin/streptomycin solution 100× (10,000 units of penicillin and 10 mg ml−1 streptomycin in 0.85% saline solution, GenClone), in a 37 °C and 5% CO2 incubator.

MCF10a and MDA-MB231 spheroids were formed starting from a culture of 20,000 cells per spheroid mixed with their medium and a 2% Matrigel (Corning Matrigel Matrix) solution in 96-well round-bottom plates (Corning Costar Ultra-Low Attachment Multiple Well Plate). After a brief centrifugation (5 min at 1,000 rpm) and 3 days of incubation (37 °C, 5% CO2), the spheroids were transferred in eight-well chambers (Cellvis) and embedded in a collagen Type-1 gel matrix (Corning Collagen I, Rat Tail, 100 mg). Media and dyes were added on top after collagen polymerization and incubated at 37 °C and 5% CO2 before imaging.

### Cell Treatments

For Glucose Starvation treatments, the spheroids were transferred in a new well of the 96-well plate containing DMEM without glucose (Gibco) for 24 hrs, before embedding in the collagen matrix.

For the Oleic Acid treatments, the spheroids were incubated with a solution of final concentration 315 μM Oleic Acid for 24 hrs, before embedding in the collagen matrix.

For Nile Red staining, the spheroids were incubated with a Nile Red (ThermoFisher) solution in DMSO (Dimethyl sulfoxide, Sigma) of final concentration of 0.3 μM for 1-2 hrs before embedding in the collagen gel matrix. Same concentration solution was added, together with media, on top of the polymerized collagen 1 hr before imaging.

### Fluorescence Imaging

The same spheroids were imaged at 3 different time points: 2 hrs after embedding, 12 hrs and 24 hrs, focusing on the plane located 100μm from the bottom of the spheroids. Data were collected with Zeiss LSM 880 inverted confocal microscope.

Fluorescence lifetime images were collected with a Zeiss LSM 880 (Carl Zeiss Microscopy) microscope equipped with a Spectra-Physics MaiTai HP laser (Spectra-Physics) for two-photon excitation, tuned at 720 nm (20% laser power), and collected using a Zeiss 10X/0.45 NA air objective. The fluorescence signal was collected by a photomultiplier tube (H7422P-40, Hamamatsu) and recorded using a FLIMbox (model A320 ISS) to obtain the lifetime information. A filter 442/46, 520/35, 484LP was used to split the signal in two channels and data collected from channel 1 were subsequently analyzed. The pixel dwell time for the acquisitions was 16 μs and the images were taken with sizes of 512 × 512 pixels. The data from each pixel were recorded and analyzed using the SimFCS software (Laboratory for Fluorescence Dynamics, University of California, Irvine, CA) to perform the phasor transformation and with custom Python code to quantify the lifetime. The phasor position was corrected for the instrument response function by acquiring the fluorescence of a solution with a known lifetime (0.4 mM Coumarin 6 dissolved in ethanol, τ = 2.5 ns). The lifetime was determined as τ = s/ωg, where τ is the fluorescence lifetime, g and s are the phasor coordinates and ω is the laser frequency (80 MHz).

Hyperspectral Images were collected using the 32 channel spectral detector module (spectral range 415-690) in Photon counting mode. The 2-Photon excitation laser was tuned to 800nm (4% laser power), the objective used was a Zeiss 10X/0.45 NA air objective, frame size is 1024×1024 pixels and to collect the data was used the sum of 16 lines. SHG images were collected tuning the 2-Photon laser at 800 nm (40% laser power)…

### Image Analysis

Image analysis was performed in MATLAB and Python. Each spheroid was segmented in user-defined regions based on the total intensity image of Nile Red. The “Core” was segmented as the center of the spheroid minus a few cell layers that appear brighter which were segmented as “Surface”. After excluding core and surface, “Invasions” were segmented by intensity thresholding to disregard the background signal coming from the collagen. The “Distance from center” was calculated as the median distance of the points at the perimeter of the mask (considering all the segmented regions) from the center of the spheroid. PLICS analysis was applied to the SHG channel. The PLICS algorithm ^50^ was applied using a 11x11 mask, yielding an image that spatially encodes the PLICS phase, a parameter that correlated with the average local size of the collagen fibers; an automatic (Otsu) threshold was applied to only consider the collagen fibers. Phasor FLIM analysis was applied to the NADH-FLIM dataset. the signal was unmixed using a 3-component model that includes free NADH (0.4 ns), bound NADH (3.2 ns) and background fluorescence with (g,s) coordinates equal to (0.344, 0.320) and (0.394,0.344) for MDA-MB-231 and MCF10A, respectively. The phasor positions for the different cell lines were calibrated using areas of collagen without cells; differences between the two are due to the different composition of the culturing media. The bound fraction of NADH was calculated as the fraction of bound NADH over the sum of the fractions of bound and free NADH. Spectral phasor analysis was applied to the Nile Red dataset.

Spectral images were transformed in spectral phasors and the spectral phase (which depends linearly from the average emission wavelength) was considered as a measure of Nile Red spectral shift. In this work, the data are shown as the median values of each parameter for each segmented region.

### Statistical Analysis

Graphs and statistical analysis (Outliers identification, One-way ANOVA/Kruskal-Wallis test) were performed using Graphpad Prism 10.2 Software.

## Acknowledgements

This study was made possible in part through access to the Laboratory for fluorescence dynamics, University of California Irvine. This work was supported by the Betty and Gordon Moore Foundation, NIH U54-CA217378 and P41GM103540.

## Author Contribution

G.T., L.S. and M.A.D. originated the idea. G.T. developed and optimized spheroids sample preparation, treatments and embedding. G.T., L.S. and F.P. performed the experiments. L.S. wrote the code for image analysis. L.S. and G.T. performed data analysis. G.T and L.S. wrote the manuscript with assistance from the other authors. All authors reviewed the manuscript.

