## Supplementary Figures for "Multimodal Phasor Approach to study breast cancer cells invasion in 3D spheroid model"

A)

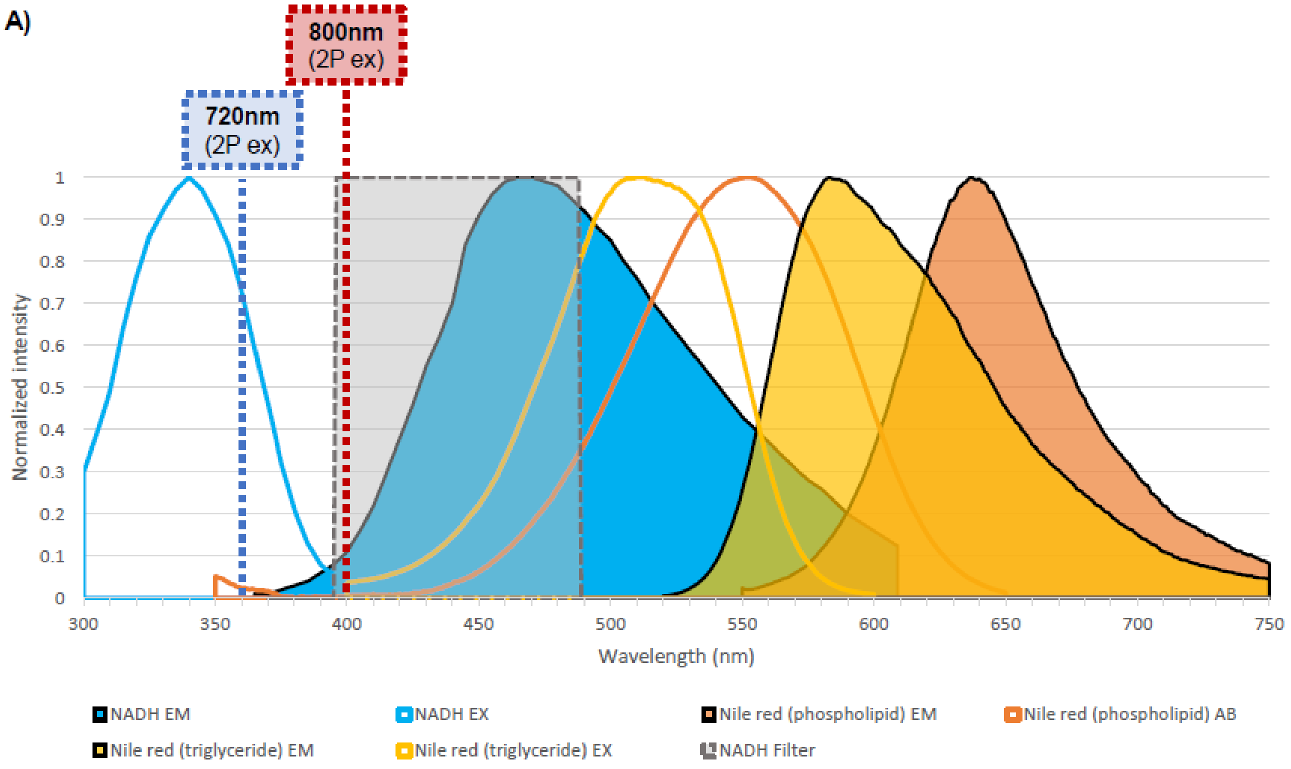


**Figure S1. Excitation and emission spectra of NADH and Nile Red. (A)** The emission filter used to collect the NADH emission is reported in grey-dotted line. Laser excitation for NADH and Nile Red are reported as dotted lines (blue, red, respectively).

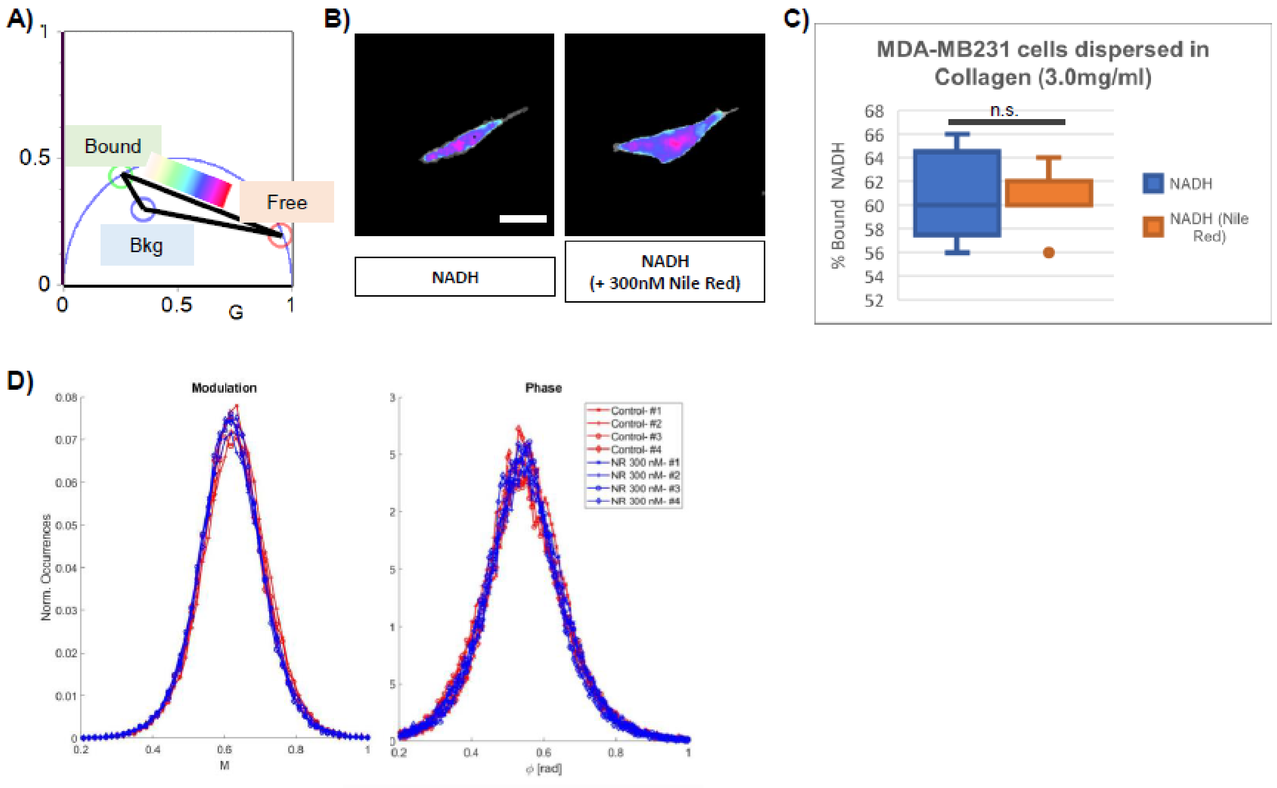


**Figure S2. FLIM NADH signature doesn't change when adding Nile Red. (A)** Lifetime Phasor Plot with metabolic trajectory between Free (red cursor) and Bound NADH (Green cursor) and background position (blue cursor). **(B)** Color map of MDA-MB231 cells embedded in Collagen I matrix label free (left) or with 300nM Nile red (right). Scale bar is 20µm. **(C)** Percentage of bound NADH is not influenced by the addition of 300nM Nile red. **(D)** Distributions of Modulation (left) and Phase (right) of NADH FLIM signal are not influenced by 300nM Nile Red (Data from MDA-MB231 3D spheroids embedded in Collagen I).

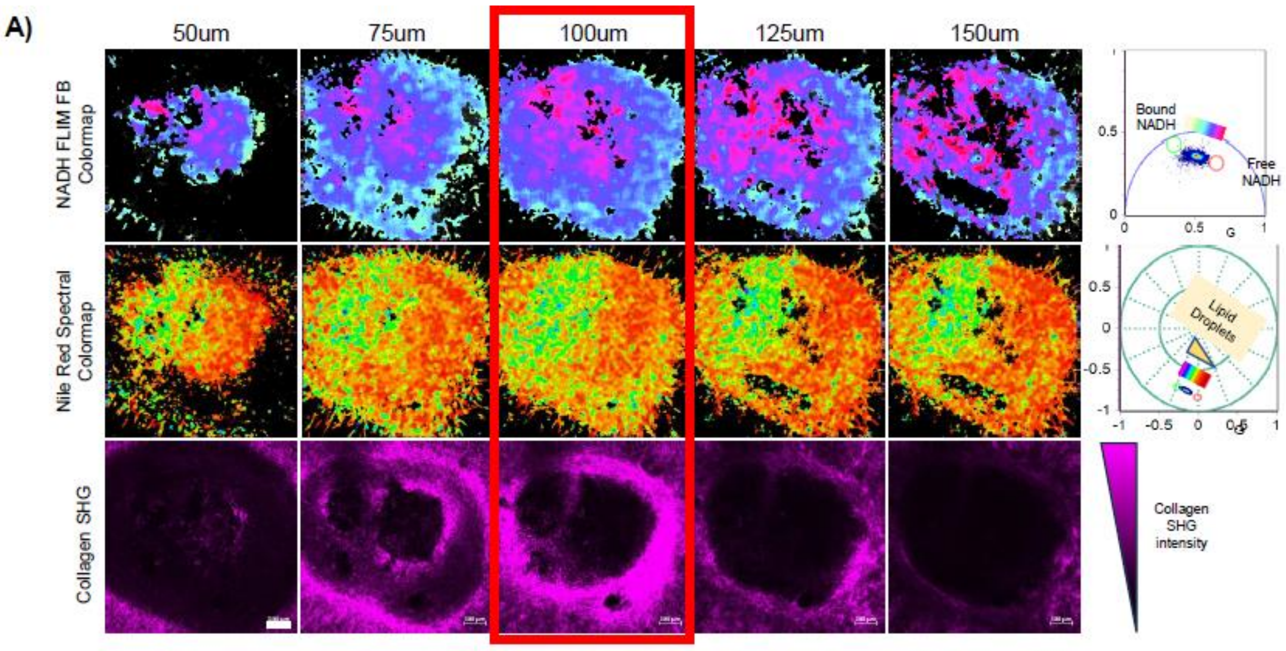


**Figure S3. Choosing a representative focal plane in the spheroids. (A)** MDA-MB231 Spheroid 24hrs after embedding in Collagen 3.0 mg/ml, different heights from the bottom of the spheroid. On the first row, NADH FLIM fraction bound signature; on the second row, Nile Red emission spectrum; on the third row, SHG intensity signal from the collagen. Scale bar is 100µm.

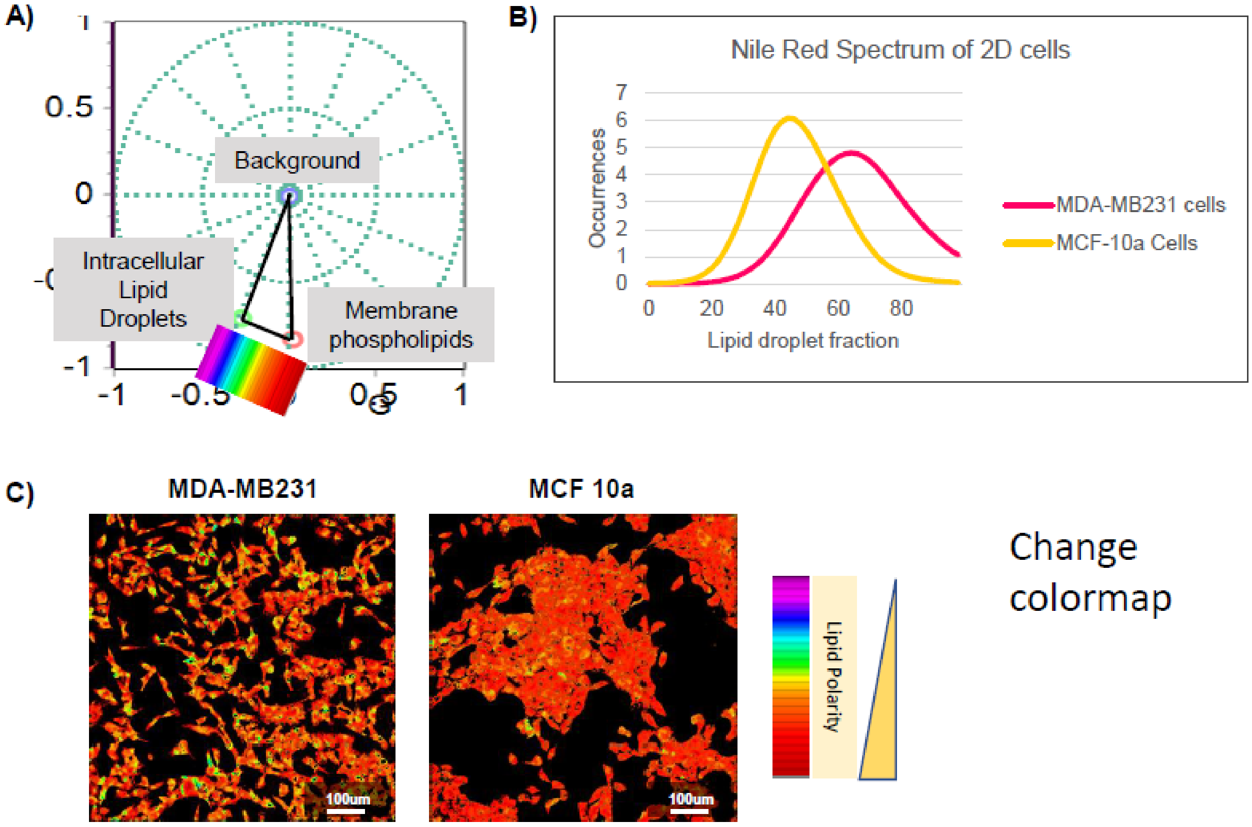


Change colormap

**Figure S4. Nile Red emission spectral signature in MDA-MB231 and MCF10a cells. (A)** Hyperspectral Phasor Plot with the position of Nile red signal for membrane phospholipids (red cursor), intracellular lipid droplets (green cursor), background (blue cursor). **(B)** Fractional contribution of the Nile red signals (membrane phospholipids, lipid droplets) for MDA-MB231 cells and MCF10a cells seeded in 2D. **(C)** Representative images of MDA-MB231 (left) and MCF10a (right) cells color coded according to the position in the phasor plot. Scale bar is 100µm.

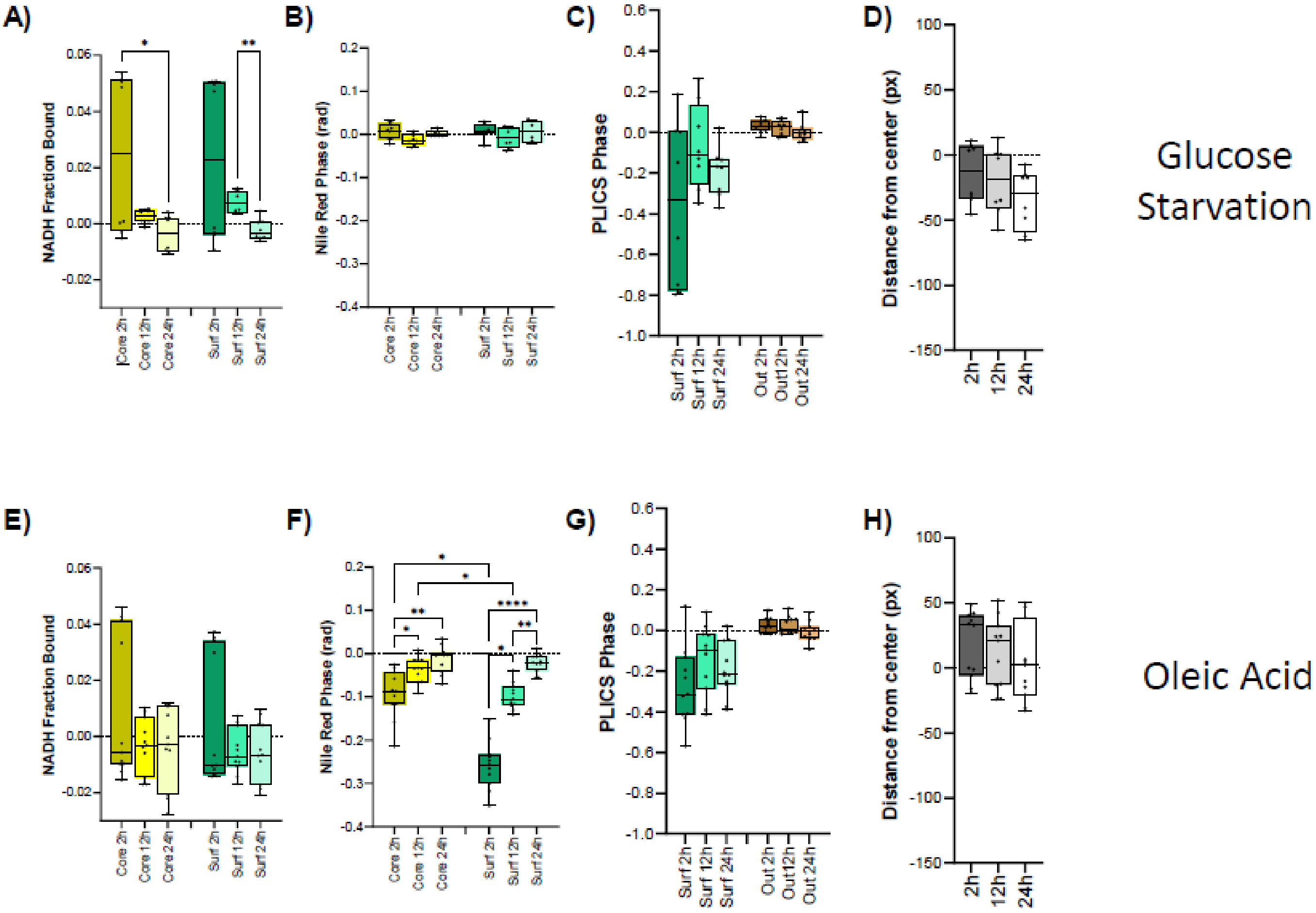


**Figure S5. Response of MCF10a spheroids to transient metabolic treatments** (Glucose Starvation A-D, Oleic Acid E-H). Metabolic FLIM response **(A, E)**, Nile Red Spectral signature **(B,F)**, Collagen remodeling **(C,G)** and distance of the perimeter from the center of the spheroid **(D,H)**. All data in panels **A-H** are shown as difference from the mean value of the control, represented by the dashed black line. Data are represented by box and whiskers plots. The box represent the median (solid line) and standard deviation. The whiskers go down to the smallest value and up to the largest. All the points are shown. One-way ANOVA (K-W test), p>0.05 (ns), 0.05>p>0.01 (*), 0.01>p>0.001 (**), 0.001>p>0.0001 (***), p<0.0001 (****).

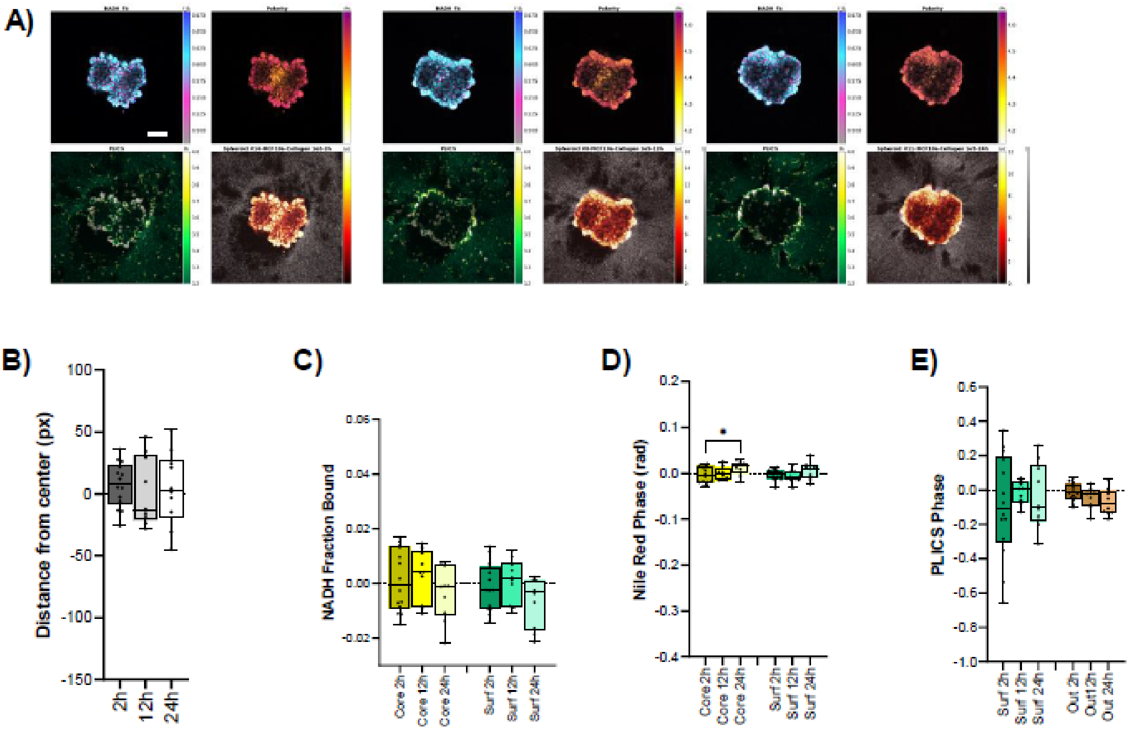


**Figure S6. Response of MCF10a spheroids to embedding in collagen with less density.** Representative images of MCF10a spheroids at 2h (A), 12h (B), 24h (C) after embedding in collagen. Scale bar is 100µm. Metabolic FLIM response (D), Nile Red Spectral signature (E), Collagen remodeling (F) and distance of the perimeter from the center of the spheroid (G). All data in panels **B-E** are shown as difference from the mean value of the control, represented by the dashed black line. Data are represented by box and whiskers plots. The box represent the median (solid line) and standard deviation. The whiskers go down to the smallest value and up to the largest. All the points are shown. One-way ANOVA (K-W test), p>0.05 (ns), 0.05>p>0.01 (*), 0.01>p>0.001 (**), 0.001>p>0.0001 (***), p<0.0001 (****).
